# Structural and functional characterisation of isolated puff adder (*B. arietans*) serine proteases

**DOI:** 10.1101/2025.06.11.659063

**Authors:** Mark C. Wilkinson, Cassandra M. Modahl, Anthony Saviola, Frank-Leonel Tianyi, Robert A. Harrison, Nicholas R. Casewell

**Affiliations:** Centre for Snakebite Research and Interventions, Liverpool School of Tropical Medicine, Liverpool, Merseyside, UK; Department of Biochemistry and Molecular Genetics, 12801 East 17th Avenue, University of Colorado Denver, Aurora, CO 80045, USA

**Keywords:** arietans, serine protease, gelatinase, venom, chromatography

## Abstract

Serine proteases are known to play a major role in the haemotoxic actions of viper venom, but compared with those from other vipers, the serine proteases of puff adder venoms have not been extensively characterised. To address this, we isolated, identified and characterised the bioactivity of the serine proteases within the venom of the Nigerian puff adder which we had previously shown to be especially rich in this class of toxin. Two distinct groups were identified, each with different protein substrate specificities. Both had similar molecular weights of 52-62 kDa, with 4-6 N-glycans, but one group consisted of trypsin-like acidic SVSPs and the other were chymotrypsin-like basic SVSPs. Each acted differently on fibrinogen: the acidic SVSPs showed thrombin-like alpha/beta-fibrinogenase activity, whereas the basic forms were shown to be alpha-fibrinogenases. The acidic SVSPs possess gelatinase activity - a novel activity for SVSPs and the first example of an SVSP acting on proteins other than those of the haemostatic system. Analysis of the transcripts of both sets of SVSPs revealed structural details of the substrate-binding sites that supported the experimental findings. The activity and sequences of the basic SVSPs show that they are very like the alpha-fibrinogenase ML-AF of *M. lebetina*, which until now was considered to be a unique SVSP. Thus, this basic SVSP and the acidic SVSP with its gelatinase activity can be considered to be atypical viper serine proteases. The gelatinase activity of the acidic SVSPs was found to vary geographically and this, alongside the regional variation in the SVMP activities that we observed previous study, is discussed with reference to the potential implications on pathology of envenoming and the development of therapeutic interventions.

## 1. Introduction

Serine proteases are found ubiquitously in the venoms of the terrestrial snake families but are especially abundant in the Viperidae (see Serrano and Maroun, 2005; Kini, 2005, for reviews). All viper SVSPs are based on a simple structural layout: a ∼26 kDa monomeric protein segment with a variable N-glycan content that results in a large range of molecular weights, from 28 to 67 kDa. From this basic structure arises a surprising variety of functions, however, all of which can disrupt one or more of the components of haemostasis. Thus, isolated viper SVSPs have been shown to have fibrinogenase activity (alpha-, beta- or less commonly alpha/beta-), fibrinolytic (via plasmin activation), platelet-activating (via PAR activation) or kallikrein-like activities (Serrano and Maroun, 2005; Kini, 2005). Most viper SVSPs are trypsin-like in their action and it is not surprising, therefore that they have also been found to activate some of the coagulation cascade clotting factors, including prothrombin and factor X (Marrakchi et al., 1995), V (Kisiel et al., 1979; Siigur et al., 1998) and VIII (Niewiarowski et al., 1979; Hill-Eubanks et al. 1989). There have been a few reports of viper SVSPs that are non-trypsin-like in their substrate specificity, but these also have also been shown to act on plasma proteins such as fibrinogen (see Discussion).

The SVSP content of the large *Bitis* sp. varies from ∼20 to 26%, and in *B. gabonica gabonica* it is the major venom component, exceeding that of the SVMP content (Calvete et al., 2007). The only study that has made a direct estimate of the SVSP content of puff adders is that of Juarez et al., (2006) quoting 19.5% for Ghanian snakes, less than that of the SVMP and CLP content. A study on Nigerian venom using the same methodology (Casewell et al., 2014, Supplement) indicates a similar level in these puff adders. The SVSP content of puff adders is clearly geographically variable, however. Thus, in a recent study which revealed substantial intraspecific variation in toxin expression in African puff adders, the level of both SVSP transcript expression and trypsin-like SVSP activity was ∼5-fold greater in Nigerian snakes than in those from Tanzania (Dawson et al. 2024). The complex nature of this Nigerian *B. arietans* SVSP complement has been highlighted by proteomic analysis (Fasoli et al., 2010). This study used 2D PAGE for a high-resolution separation of the venom proteins for identification via mass spectrometric analysis and identified multiple SVSP proteo- and glycoforms, grouped into 5-6 glycosylation trains with an overall range of isolectric points between 4 and 7.5, and molecular weights of 37 to 60 kDa.

Previous studies using traditional chromatographic methods to isolate and characterise individual puff adder SVSPs have been confined to those responsible for kallikrein-like (kinin-releasing) activity (Sekoguchi et al., 1986; Nikai et al., 1993 Megale et al., 2018). Such SVSPs were isolated and found to possess a wide range of molecular weights, from 33 to 58 kDa, due to differences in the degree of N-glycosylation, as highlighted above. Considering the established functional diversity of SVSPs and the demonstrated complexity and content of the puff adder SVSPs, it is unlikely that kinin-releasing activity is their only role in puff adder venom, however. All the aforementioned studies have provided some insight into the role of SVSPs in the toxicity of puff adder venom but a complete picture of the SVSPs and their precise bioactivities is required to fully understand the diverse pathological effects of the venom and identify potential therapeutic targets. Here, we describe the purification of, and the structural and functional characterisation of puff adder SVSPs. The study was prompted by our observation of pronounced differences in PMSF-inhibited gelatinase activities of venom from Nigerian and Tanzanian snakes (Wilkinson et al. 2025). The bulk of the work described here focusses on the serine proteases in venoms from the Nigerian snakes, since a related transcriptome was available that enabled a full analysis of the variety of SVSPs and their functions.

## 2. Materials and methods

### 2.1. Venom

Venoms were extracted from five Nigerian (NGA) and four Tanzanian (TZA) *B. arietans* specimens maintained within the herpetarium at the Centre for Snakebite Research and Interventions at the Liverpool School of Tropical Medicine (LSTM). Venoms from individual specimens were labelled, immediately frozen at -20°C and later lyophilised for long term storage at 2 – 8°C. To deliver the volume of venom required for the multiple analyses described below, venom extractions were performed on multiple occasions from each snake. The other venoms used in this study (for the gelatin zymograms, Fig S1) were taken from stocks held at LSTM; these are either from snakes previously kept in the herpetarium (Ghanaian and Nigerian) or samples taken from wild-caught animals over many years and supplied to LSTM.

### 2.2. Reagents

All reagents used were of analytical reagent grade and, unless otherwise specified, were purchased from Merck Life Science, Watford, UK or Fisher Scientific, Loughborough, UK.

### 2.3. Serine protease purification

All chromatography was carried out using either an AKTA LC system (Cytiva) or a Vanquish HPLC system (Thermo Fisher). Chromatography buffers were freshly prepared and vacuum filtered (0.1 um) immediately prior to use. The initial step in the isolation procedure was a separation of whole venom on a size exclusion chromatography (SEC) column. Twenty mg of freeze-dried venom was resuspended in 1.5 mL ice-cold PB5.2 (50 mM sodium phosphate pH 5.2) and centrifuged at 10,000 xg for 10 mins. The supernatant was immediately loaded onto a 120 mL column of Superdex 200HR equilibrated in PB5.2. The column was operated at a flow rate of 1.0 mL/min and 2 mL fractions were collected after the void volume. Elution was monitored at 214 and 280 nm. SDS-PAGE analysis and protease assays were carried out on all protein-containing fractions to determine which to select for the second stage of isolation. For all venoms, serine protease (SVSP) activity, indicated by PMSF-inhibited casein degradation found in the first main peak of the SEC separation, with bands visible on SDS-PAGE in the 50-60 kDa range (reducing conditions). These were pooled and applied to a 1 mL HiRes Capto S column equilibrated in PB5.2. The unbound material in the flow-though (acidic SVSPs) was retained and then elution of the bound (basic) SVSPs was carried using a 25-column volume (CV) gradient of 0 - 0.70 M NaCl in PB5.2. The flow rate was 0.6 mL/min. The acidic SVSPs in the unbound fraction were immediately buffer exchanged into 50 mM Tris-Cl, pH 8.5 using a 50 mL Sephadex G25 desalting column. These were then loaded onto a 1 mL Mono Q column equilibrated in the same buffer. The acidic SVSPs were eluted from the column using a 25 CV gradient of 0-0.4 M NaCl in 50 mM Tris-Cl, pH 8.5. The column was operated at 0.6 mL/min and 0.5 mL fractions were collected. Elution was monitored at 214 and 280 nm. The final purification step on reverse-phase HPLC (RP-HPLC) was performed using a Biobasic C4 column (2.1 x 150 mm, Thermo Fisher). The flow rate was 0.2 mL/min and proteins were separated in the following gradient of acetonitrile in 0.1% trifluoroacetic acid; 0-32%/5 mins; 32-44%/30 mins; 44-70%/2 mins. Elution was monitored at 214 nm.

### 2.4. Analytical methods

#### 2.4.1 SDS-PAGE

Immediately prior to this analysis, the venoms were reconstituted in PBS (50 mM sodium phosphate, 0.15 M NaCl pH 7.2) and maintained on ice. Samples were prepared for reducing SDS-PAGE by adding sample buffer to a final concentration of 2% SDS, 5% β-mercaptoethanol and then heating at 85°C for 5 mins. For non-reducing gels the heat treatment was omitted. Electrophoresis was performed on 4-20% acrylamide gels (BioRad TGX) using a Tris-glycine buffer system, followed by staining with Coomassie Blue R250. The molecular weight markers used were Thermo-Fisher PageRuler or Promega Broad Range (see figure legends).

#### 2.4.2. Deglycosylation of isolated proteins

In preparation for deglycosylation, a 20 μL sample was denatured by heating for 5 mins at 85°C following the addition of 2.0 μL 1% SDS and 0.5 μL 1 M DTT. After cooling to room temperature,1.7 μL of 10% NP-40 was added and then 0.5 μL PNGase F (at the supplied concentration) was added to start the reaction. Incubation was carried out at 42°C for 3 hours or overnight at room temperature. Where deglycosylation was carried out under native conditions for gelatin zymograms (Fig. S5), the denaturing step was omitted and deglycosylation was performed at 37°C.

#### 2.4.3. General protease assays

To distinguish serine protease activities from those of the metalloprotease the ability of the purified proteases to degrade casein and insulin B was determined in the presence of EDTA (metalloprotease inhibitor) or PMSF (serine protease inhibitor). Insulin B degradation was also used to give an indication of the degree of specificity of action of the protease as evidenced by the number of peptide products observed on HPLC following digestion. Digestion of angiotensin I was tested because of the well-documented hypotensive action of puff adder venoms, but also to provide further information on peptide bond specificities. Chromogenic substrates, each with a different amino acid in the key position, were used to determine the substrate specificity of the SVSPs allowing their classification as trypsin-, chymotrypsin- or elastase-like. The SVMPs were also tested against the fluorogenic substrate ES010 since this is routinely used in our laboratory for high-throughput work.

##### 2.4.3.1 Casein degradation with SDS-PAGE analysis

Venom or protein samples were incubated in TBSC (35 mM Tris-Cl, 0.5 M NaCl, 1 mM CaCl_2_, pH 7.4) with beta casein at a ratio of 30:1 (w/w) casein:sample protein; using a final concentration of 1.0 mg/mL casein. Incubation was carried out at 37°C for 2 hours. SDS-PAGE sample buffer was added (final concentration 2% SDS, 5% β-mercaptoethanol) to stop the reaction. This was then heated at 85°C for 5 mins. The extent of casein degradation was assessed using SDS-PAGE (see 2.3.5.1). In EDTA inhibition experiments, EDTA was added to the protein sample to a final concentration of 5 mM and incubated for 15 mins prior to the addition of casein to start the reaction. Where PMSF was used as an inhibitor, this was added to a final concentration of 2 mM from a 20 mM stock solution in freshly prepared in methanol. This pre-incubation was carried out for 60 mins (room temperature) prior to addition of casein.

##### 2.4.3.2. Insulin B or angiotensin I digestion with RP-HPLC analysis

Venom or protein samples were incubated in TBSC with insulin B chain or angiotensin I at a ratio of 30:1 (w/w) insulin B:sample protein, using a final concentration of 0.02 mg/mL insulin B/ angiotensin I. Where inhibitors were used (insulin B assay only), these were pre-incubated with the protein sample as in section 2.3.4.1. Incubation was carried out at 37°C for 90 mins and the reaction was stopped by the addition of trifluoracetic acid (TFA) to 1%. An aliquot containing the equivalent of 1 μg insulin B/ angiotensin I was analysed by RP-HPLC using a Biobasic C4 column (2.1 x 150 mm). The flow rate was 0.3 mL/min and the separation was carried with the following gradient of acetonitrile in 0.1% trifluoroacetic acid; 0-36%/40 mins; 36-70%/3 mins. Elution was monitored at 214 and 280 nm.

##### 2.4.3.3. Chromogenic substrate assays

Assays using the chromogenic substrates BAEE (N-alpha-benzoyl-L-arginine ethyl ester) and BTEE (N-benzoyl-L-tyrosine ethyl ester) were carried out in TBSC with the substrate at a concentration of 0.25 mM and the protease at a final concentration of 2-5 μg/mL. The reaction was followed in quartz cuvettes at 253 nm (BAEE) or 256 nm (BTEE). Where nitroanilide esters (N-succinyl-A-A-A-p-NA, N-methoxysuccinyl-A-A-P-V p-NA, N-succinyl-A-A-P-L p-NA, N-succinyl-A-A-P-F p-NA) were used the final concentrations were 0.2 mM substrate in TBSC and 2-5 μg/mL protease. The reaction was followed at 410 nm. In all cases specific activity was calculated as μmol product formed per minute per mg protease. The mM extinction coefficients used for this calculation were BAEE, 1.070 (253 nm); BTEE, 0.964 (256 nm) and for all the nitroanilides, 8.800 (410 nm).

#### 2.4.4. Functional protease assays

Four assays were carried out to determine the abilities of the SVSPs to degrade proteins that may be of functional and pathological significance. Degradation of gelatin (predominantly collagen I) or laminin-rich basement membrane material are possible indicators of the ability to cause tissue damage. Potential coagulopathic activities were measured by the ability to degrade the key coagulation factors, fibrinogen and prothrombin. In the latter case, the specific proteolytic action to generate thrombin can be determined.

##### 2.4.4.1 Gelatin zymogram assay

The gels (10% acrylamide) for this assay were prepared using the method of Laemlli (1970) with the adaptations of Fling and Gregerson (1986). Gelatin was prepared by briefly heating a 20 mg/mL solution at 50°C, then adding this to the gel mix immediately prior to casting, such that the final gelatin concentration was 2 mg/mL (0.2% w/w). Where inhibitors were tested, these were pre-incubated with the protein sample as in section 2.4.3.1. Venom or protein samples were then prepared for the zymogram assay by adding standard SDS-PAGE sample buffer, but with no reductant and a final SDS concentration of 1.6% (w/v). These were not heated. Following electrophoresis, the method of (Toth et al., 2012) was used to visualise gelatinase activity. Thus, the gel was rinsed, with shaking, in 2.5% (v/v) Triton X-100 for 30 mins, then washed (3 x 10 mins) with distilled water to remove the detergent. The buffer used for development was 50 mM Tris-Cl, 200 mM NaCl, 5 mM CaCl_2_, 0.02 % (v/v) Brij 100, pH 7.8. The gel was washed for 10 mins in this buffer, then this was replaced with fresh development buffer and incubated overnight at 37°C. To visualise gelatin degradation zones, the gel was stained for 60 mins with Coomassie Blue R250 and destained until clear bands were visible against a blue background.

##### 2.4.4.2 Digestion of basement membrane proteins

Basement membrane material (Geltrex, Thermo Fisher) was diluted with PBS to 10% of its supplied concentration (10-18 mg/mL) and stored in aliquots at -20°C. For the assay, a fresh aliquot was carefully thawed out and immediately placed on ice. Digestions were set up with basement membrane material:protein sample at 30:1 (w/w) and performed at 37°C for various time periods between 2 mins and 4 hours. Following this, SDS-PAGE sample buffer was added and the sample heated at 85°C to stop the reaction. The extent of basement membrane protein degradation was then assessed using SDS-PAGE (see section 2.4.1).

##### 2.4.4.3 Digestion of plasma proteins (prothrombin and fibrinogen.)

In both the prothrombin and fibrinogen degradation assays, these two substrate proteins were used in the assay at a final concentration of 1.0 mg/mL in TBSC and venom/pure protein samples were added at a ratio of 30:1 substrate:protein sample to start the reaction. Incubation was carried out at 37°C for 60 mins. Following this, SDS-PAGE sample buffer was added, the sample heated at 85°C to stop the reaction and the extent of substrate protein degradation was then assessed using SDS-PAGE (see section 2.4.1). An aliquot equivalent to 1.0 μg of the substrate protein was loaded per lane of the gel.

### 2.5 Trypsin digestion and MS/MS analysis

In preparation for this, the relevant proteins were desalted on a RP-HPLC column, as above. These were then dried in a centrifugal evaporator, 20 μL H_2_O was added and then re-dried. These proteins were then prepared for LC-MS/MS and the resulting fragmentation spectra were searched against an in-house Nigerian and Tanzanian *B. arietans* venom gland derived protein sequence database according to the methods in Wilkinson et al., (2025).

## 3. Results

### 3.1. Gelatinase assays

Whilst studying the activities of the prominent SVMPs (arilysins) of puff adder venoms (Wilkinson et al. 2025), we observed a strong gelatinolytic activity in many venoms that was shown to be due to serine protease (SVSP) activity (Fig. 1A). The figure shows an example of two of each of venoms from the five Nigerian and four Tanzanian snakes kept at LSTM. The NGA venoms contained a strong gelatinase activity at around 55-60 kDa which was most active in NGA snake 011. No such activity was observed in any of our TZA venoms, not just the two shown here (see Fig. S1). The pronounced difference between the activity in the Nigerian and Tanzanian venoms seen in Fig.1 is typical of the geographical variation observed using a wide range of puff adder venoms from stocks held at LSTM, which can be seen in Fig. S1. In these examples, venom from Eswatini and Kenyan snakes held at LSTM showed gelatinase activity as did four stock Nigerian venoms, and stock venoms from Ghana, Malawi, Zimbabwe and one South African venom. A second sample of South African venom and one from Saudia Arabia showed no gelatinase activity. Zymogram B shows the variation in gelatinase activity in venom from three different Kenyan puff adders held at LSTM: not only in the degree of activity, but differences in the levels of two different forms of the protein can also be seen.

**Fig. 1.**
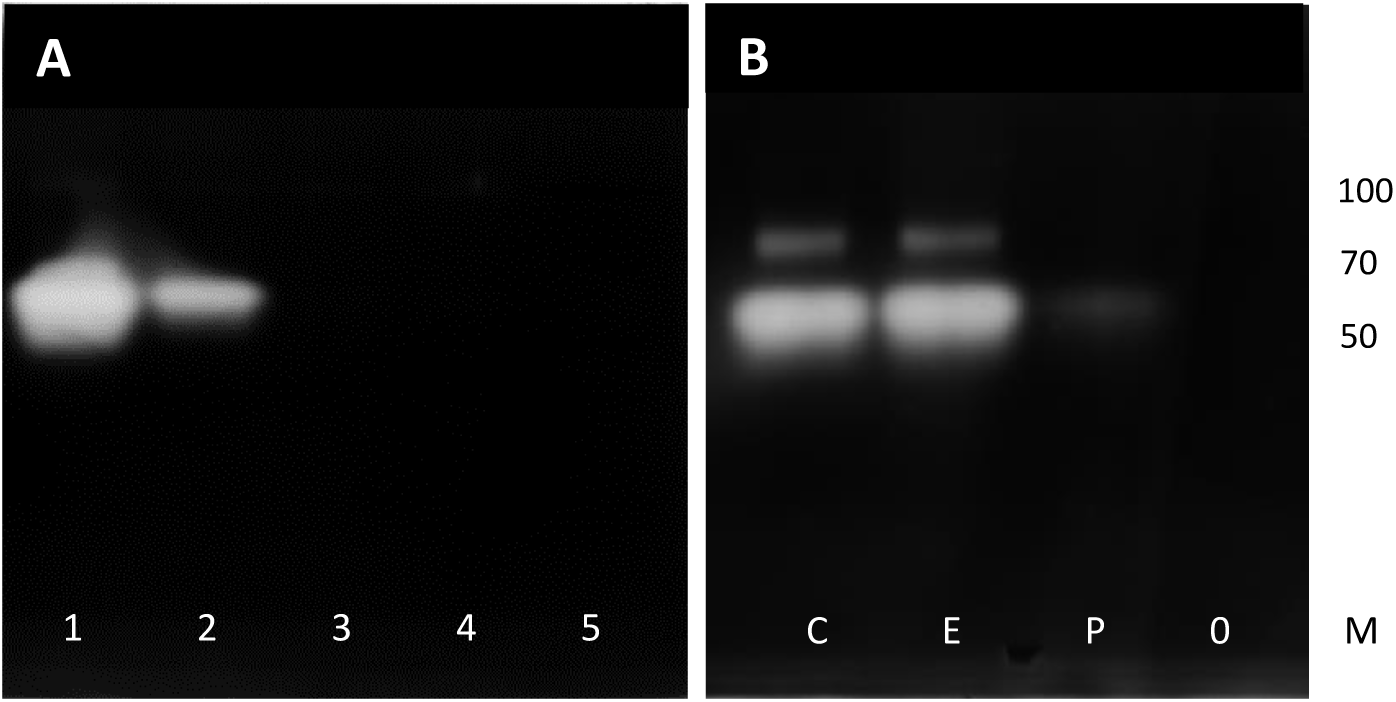
Gelatinase activity (gelatin zymogram) of Nigerian and Tanzanian *B. arietans* venoms. The zymograms were prepared with 0.2% (w/w) gelatin in a 10% acrylamide SDS-PAGE gel. The gels were run under non-reducing conditions and 2 μg of venom was loaded per lane. The zymograms were then incubated as in section 2.4.4.1 and then visualised by staining with Coomassie Blue R250. **A:** Zymogram of venoms: lanes 1, 2: NGA snakes 011 and 016; lanes 3, 4: TZA snakes 005 and 006; lane 5: control (no venom). **B:** Zymogram of NGA 011 venom treated with protease inhibitors. The venoms were pre-incubated with either EDTA or PMSF as described in the main text and then run as for zymogram A. Lanes C, controls (un-inhibited venoms); E, EDTA-treated venoms; P, PMSF-treated venoms; 0, no venom; M, markers as in zymogram A. The molecular weight markers (lane M, Thermo Page Ruler, indicated in kDa,) were run under reducing conditions and visualised after fully destaining the gel.

The same assay format was used to measure the effects of protease inhibitors in order to determine the class of the protease responsible for the activity. EDTA or PMSF, inhibitors of SVMPs and SVSPs, respectively, were incubated with the most gelatinolytic NGA venom (Fig 1B). The activity in thus venom was inhibited by PMSF, but not EDTA, and therefore the 55-60 kDa gelatinases are SVSPs. Since this activity is more commonly found to be due to SVMPs in vipers, we set about to isolate and characterise the SVSP responsible for the gelatinase activity. The work was focussed on the Nigerian and Tanzanian venoms because a full transcriptome was available for snakes from these two regions (Dawson et al. 2024).

### 3.2. Isolation and biochemical characterisation of the serine proteases (SVSPs)

Following SEC of NGA venoms, the second elution peak (Fig. S2 peak 2) contained multiple proteins in the 50-65 kDa size range and was rich in protease activity. Neither EDTA or PMSF could fully inhibit this activity, indicating that there are both SVSPs and SVMPs in this peak. The SVSP gelatinase activity observed in the whole NGA venom (Fig. 1) was also found here, so this venom was used to develop isolation methods for the individual proteases in the 50-65 kDa size range. The material from this peak was applied to a cation exchange chromatography column (see Sec. 2.3). This resulted in a large amount of material passing through unbound as well as material which bound and then eluted in the NaCl gradient in multiple peaks (Fig. 2A). Peaks 2 and 3 were found to contain serine protease activity, inhibited by PMSF but not EDTA, using both casein and insulin B degradation assays (see Fig. S3 and S4 for the results for peak 3). Peak 2 contained a single 52 kDa protein and peak 3 contained proteins at 52 and 56 kDa (Fig. 2B). RP-HPLC was used to separate the individual SVSPs and, following removal of the solvent and resuspension in PBS, their protease activity was found to have been retained. This RP-HPLC step enabled a full separation of the two proteins in cation exchange chromatography peak 3 (Fig. 2D, peak 1 is 56 kDa; peak 2 is 52 kDa). These two proteins and that from peak 2 will be referred to henceforth as basic SVSPs.

**Fig. 2.**
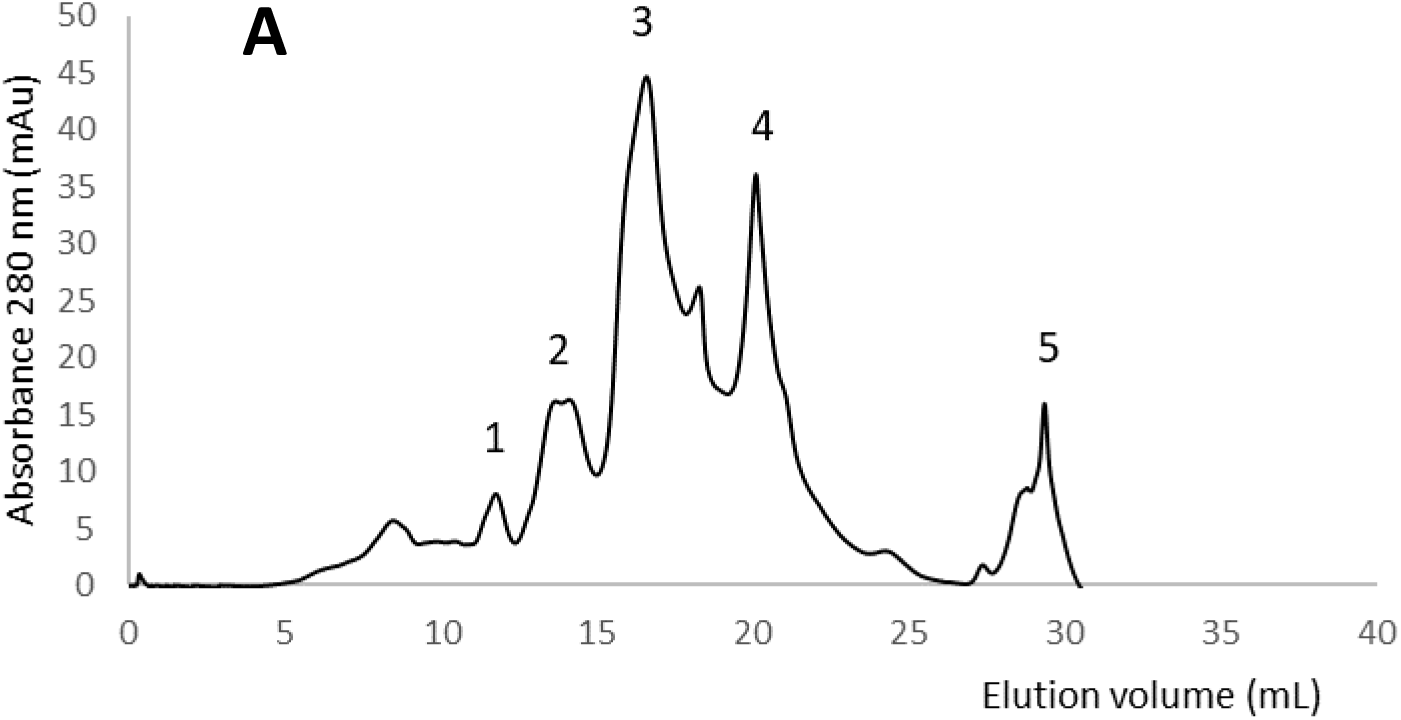
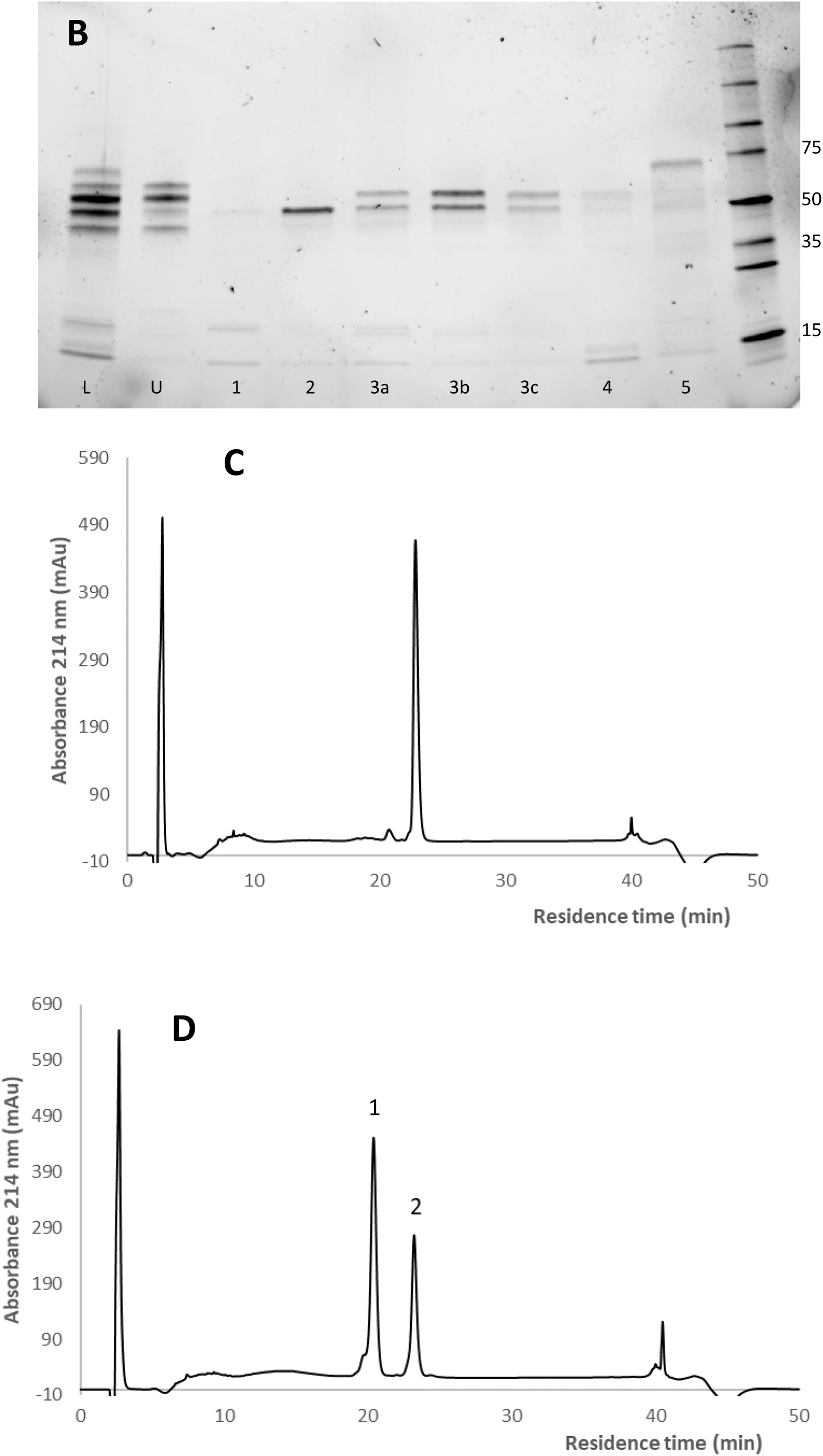
Purification of the basic SVSPs from NGA SEC peak 2. **A:** The proteins in SEC peak 2 (Fig. S2) were applied to a 1 mL HiRes Capto S column equilibrated in 50 mM sodium phosphate pH 5.2. Elution was carried using a gradient of 0 - 0.7 M NaCl and monitored at 280 nm. The proteins were loaded onto the column independently of the program used to perform the chromatography, so the peak of unbound protein used for the subsequent anion exchange chromatography (Fig. 3) is not visible here**. B:** SDS-PAGE of the main fractions from the cation exchange run. L, load; U, unbound proteins; 1-5 fractions from the respective chromatography peaks (peak 3 was run as three fractions a-c); M, markers (Promega Broad Range), molecular weights of key markers are indicated in kDa. The gel was 4-20% acrylamide, stain-free (BioRad). **C** and **D:** C4 RP-HPLC (see Sec. 2.3) of the main proteins from cation exchange chromatography, C, peak 2; D, peak 3b.

The proteins that did not bind to the cation exchange column (see lane U, Fig. 2B) also possessed SVSP activity. These were dialysed against 50 mM Tris-Cl, pH 8.5 and subjected to anion exchange chromatography. The bulk of the bound protein eluted in one main peak (peak 3, Fig. 3) which contained strong bands at 58 and 62 kDa. An earlier-eluting peak (1) contained a band at 43 kDa. Using a casein assay, the proteins in peak 3 were shown to contain only serine protease activity, inhibited by PMSF but not EDTA (Figs. S3 and S4). They were not as active towards casein as were the basic SVSPs and, in stark contrast to the latter, the acidic SVSPs were unable to cleave any of the peptide bonds of insulin B (Fig. S4). The proteins in peaks 2 and 3 were subject to RP-HPLC (Fig. 3C) and resulted in a good separation of the two proteins in cation exchange chromatography peak 3 (Fig. 3C, peak 1 is 60 kDa; peak 2 is 56 kDa). As in the case of the basic SVSPs, both retained their protease activity following this final isolation step.

**Fig. 3.**
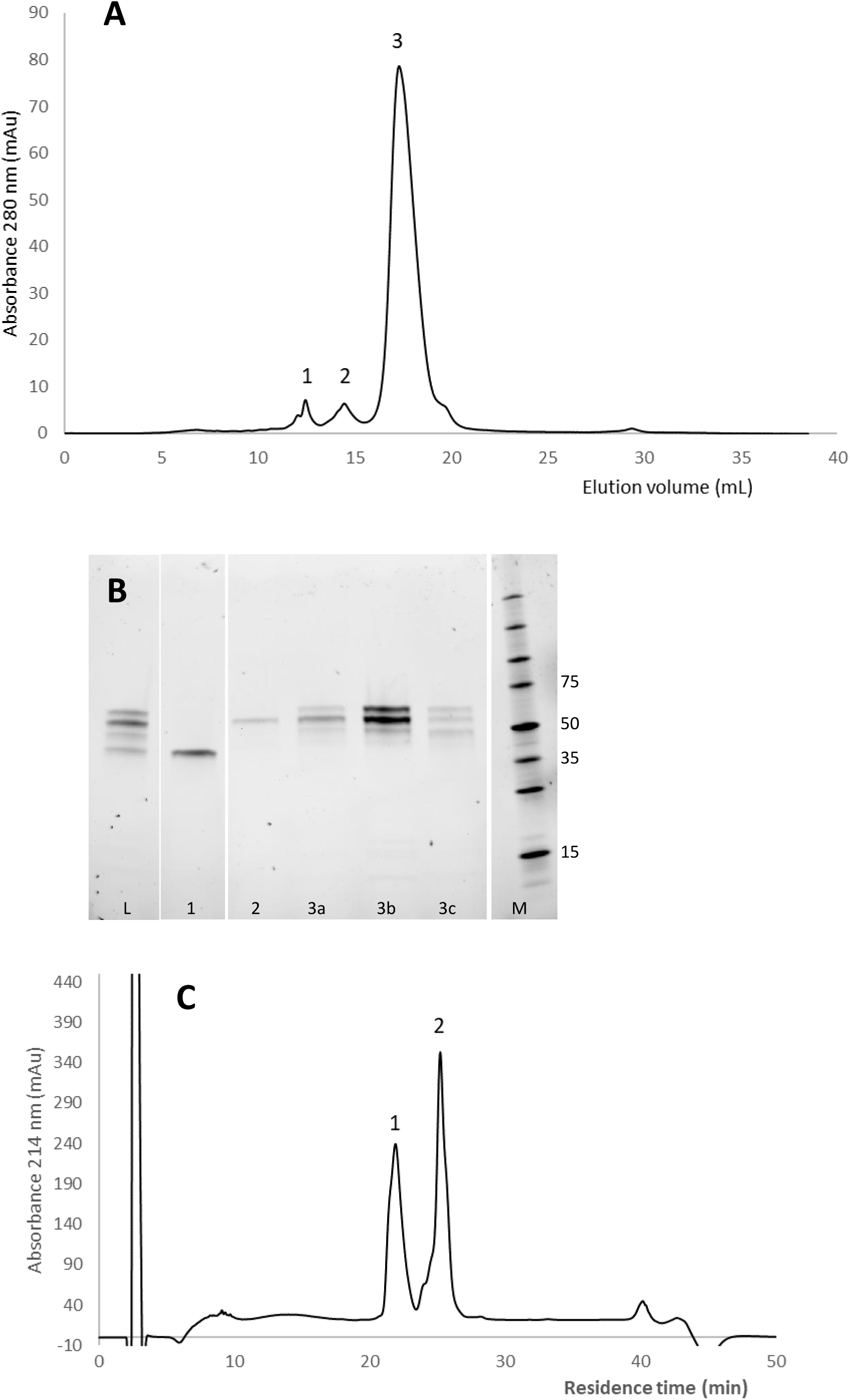
Purification of the acidic SVSPs from NGA SEC peak 2. **A**: The unbound material from the cation exchange step (Fig. 2) was dialysed against 50 mM Tris-Cl, pH 8.5 and loaded onto a 1 mL Mono Q column equilibrated in the same buffer. The acidic SVSPs were eluted from the column using a gradient of 0-0.4 M NaCl and elution was monitored at 280 nm. **B**: SDS-PAGE of the main fractions from the cation exchange run. L, load; 1-3, fractions from the relevant chromatography peaks (peak 3 run as three fractions a-c); M, markers (Promega Broad Range), molecular weights of key markers are indicated in kDa. The gel was 4-20% acrylamide, stain-free (BioRad). **C:** C4 RP-HPLC (see Sec. 2.3) of the main protein PNGase F treatment was used to determine the number of N-glycans on the basic and acidic SVSPs (Fig. 4). The purified proteins in the main ion exchange peaks were used for this analysis. In both cases, glycan cleavage was incomplete, but this allowed visualisation of the intermediates with the full range of N-glycans. For both proteins, the lowest molecular weight band was around 24 kDa, a size that would be predicted from the sequence (see Discussion). From there upwards (increasing molecular weight) and compiling the results from all 4 lanes on the gel, bands can be seen in 2-3 kDa steps which would account for the protein chain plus 1 (just above the PNGase F band at ∼30 kDa), 2, 3, 4 (alongside the 50 kDa marker), 5 and 6 N-glycans. This would suggest that the two bands in the basic SVSP peak have 4 (the 52 kDa form) and 5 (the 56 kDa form) N-glycans, and that of acidic SVSPs, 5 (the 56 kDa form) and 6 (the 60 kDa form) N-glycans.

**Fig. 4.**
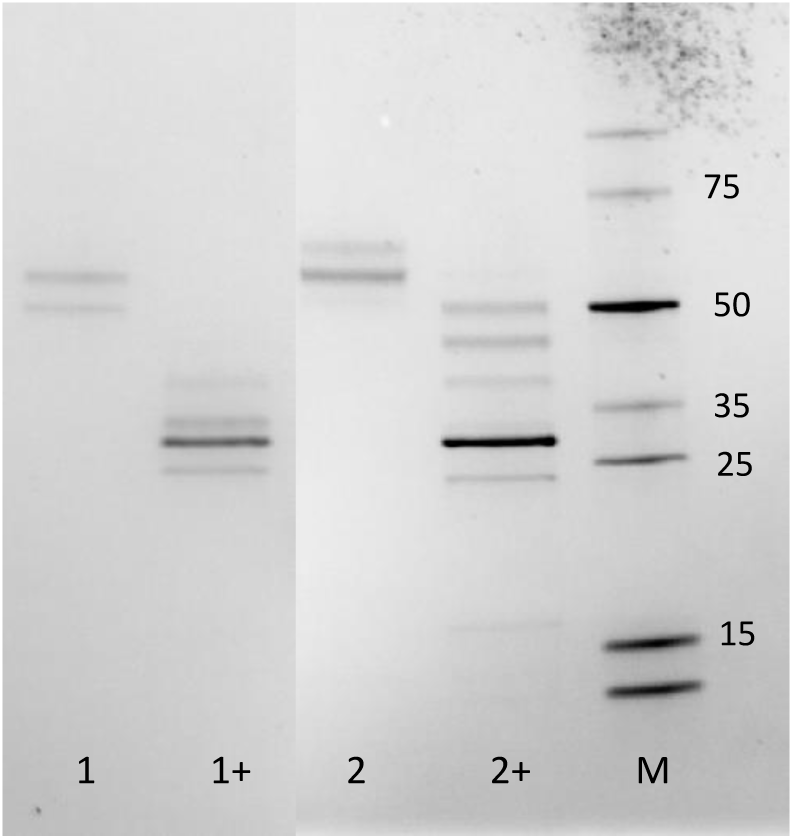
De-glycosylation of the pure SVSPs. Samples were denatured then treated with PNGase F for 3 hours at 42°C. Lanes 1, 1+, basic SVSPs; lanes 2, 2+, acidic SVSPs. Lanes indicated + are the PNGase F-treated proteins. The strong band at around 30 kDa in these lanes is PNGase F. The molecular weights of the markers (Promega Broad Range) are indicated in kDa on the right. The strongly-stained band at ∼30 kDa is the PNGase F.

A set of chromogenic esterase substrates were used to determine the substrate site (P_I_) specificities of the purified serine proteases (Table 1). The acidic SVSPs showed activity against just the R-containing BAEE substrate, confirming them to be trypsin-like SVSPs. The basic SVSPs had no activity against BAEE, nor towards the BTEE substrate traditionally used to determine chymotrypsin-like SVSP activity. Using a set of nitroanilide esters with Val, Leu, Ala and Phe in the key P_I_ site, the basic SVSPs showed a small but definite activity towards that containing phenylalanine (N-succinyl-A-A-P-F-pNA).

**Table. 1.** Determination of the primary substrate specificity of NGA SVSPs.

| | Specific activity ($\mu\text{mol product formed/min/mg SVSP}$) | | | | | |
| --- | --- | --- | --- | --- | --- | --- |
|  | Ethyl esters |  | Nitroanilide esters |  |  |  |
| **Enzyme** | **BAEE** | **BTEE** | **Ala** | **Leu** | **Val** | **Phe** |
| Trypsin | 36.5 | ND | ND | ND | ND | 0 |
| Chymotrypsin | ND | 27.3 | 0 | 1.85 | 0 | 12.2 |
| Elastase | ND | ND | 2.50 | 8.56 | 1.83 | 0 |
| Acidic SVSP | 7.13 | 0 | 0 | 0 | 0 | 0 |
| Basic SVSP | 0 | 0 | 0 | 0 | 0 | 0.031 |

The RP-HPLC purified SVSPs were digested with trypsin and subjected to MS/MS. The resulting data was searched against a list of 24 SVSPs sequences identified in the transcriptome of an NGA snake (Dawson et al. 2024). The best matches are shown in Fig. 5. The predicted pI and number of N-glycans matches well, respectively, with the chromatographic behaviour and the number of N-glycans determined experimentally. Although there is much homology between the two forms, there are critical differences in the substrate binding sites (see Discussion).

**Fig. 5.**
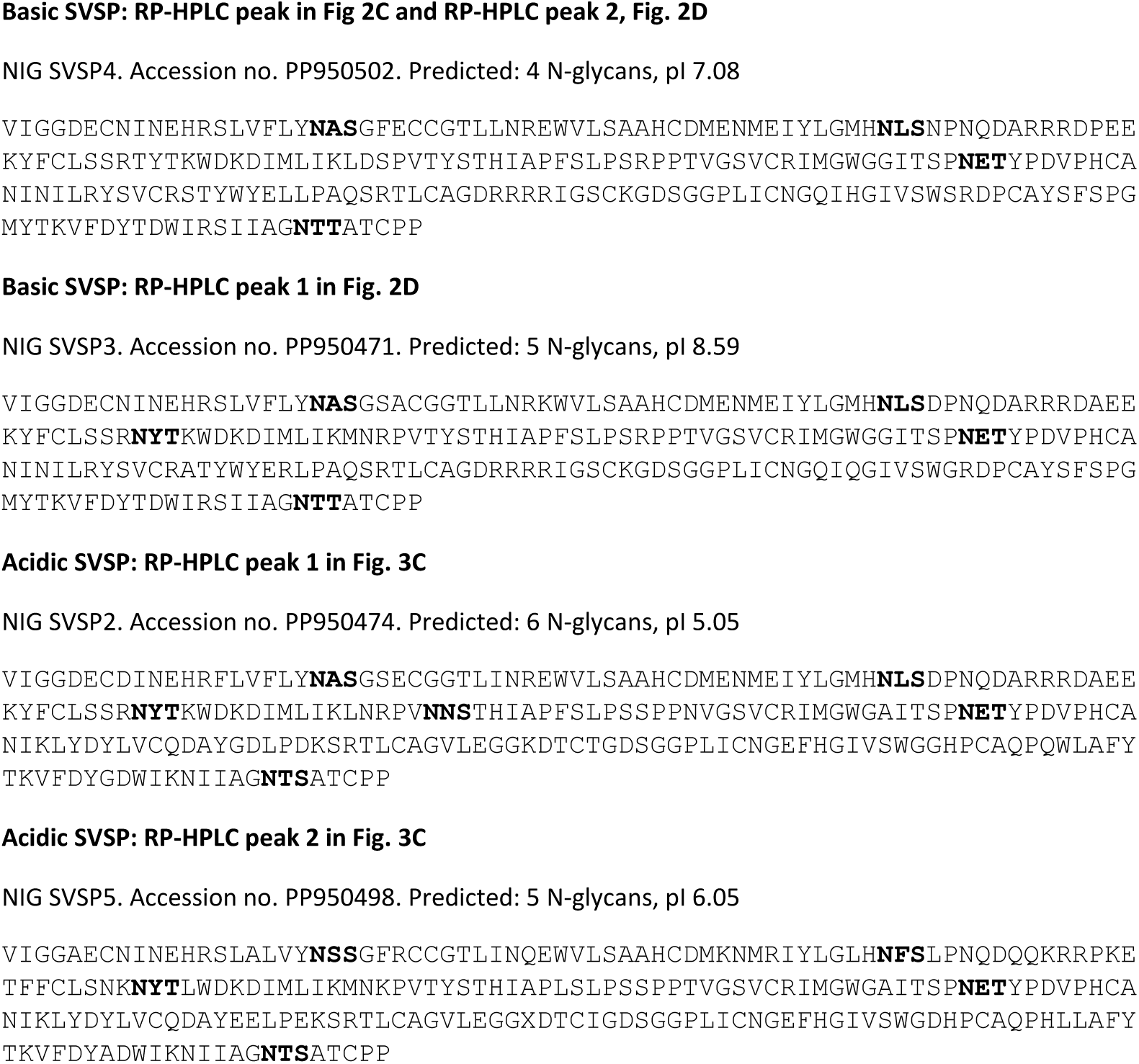
Amino acid sequences of the transcripts matching to the SVSPs isolated from NGA venoms. Ten μg of the pure proteins was digested with trypsin and the resulting peptide mix was analysed by LC-MS/MS (Sec. 2.5). Fragmentation spectra were searched against an in-house *B. arietans* (NGA) venom gland derived protein sequence database using Mascot. Predicted N-glycan sites in the sequence are indicated in bold font.

### 3.3. Activity of the isolated proteases in the functional assays

#### 3.3.1 Gelatinase activity

The studies using whole venom in gelatin zymogram assays showed that the gelatinase activity in the NGA venoms was due to the action of serine proteases (see Fig. 1). Using the purified NGA SVSPs in the same assay it became clear that this activity resides solely with the acidic SVSPs (Fig. 6A). The activity of the latter was not inhibited by EDTA, but was by PMSF. The basic SVSPs demonstrated no gelatinase activity. Both RP-HPLC purified 56 and 60 kDa acidic SVSPs were active (Fig. 6B), with gelatin-degrading activity observed at their respective molecular weights.

**Fig. 6.**
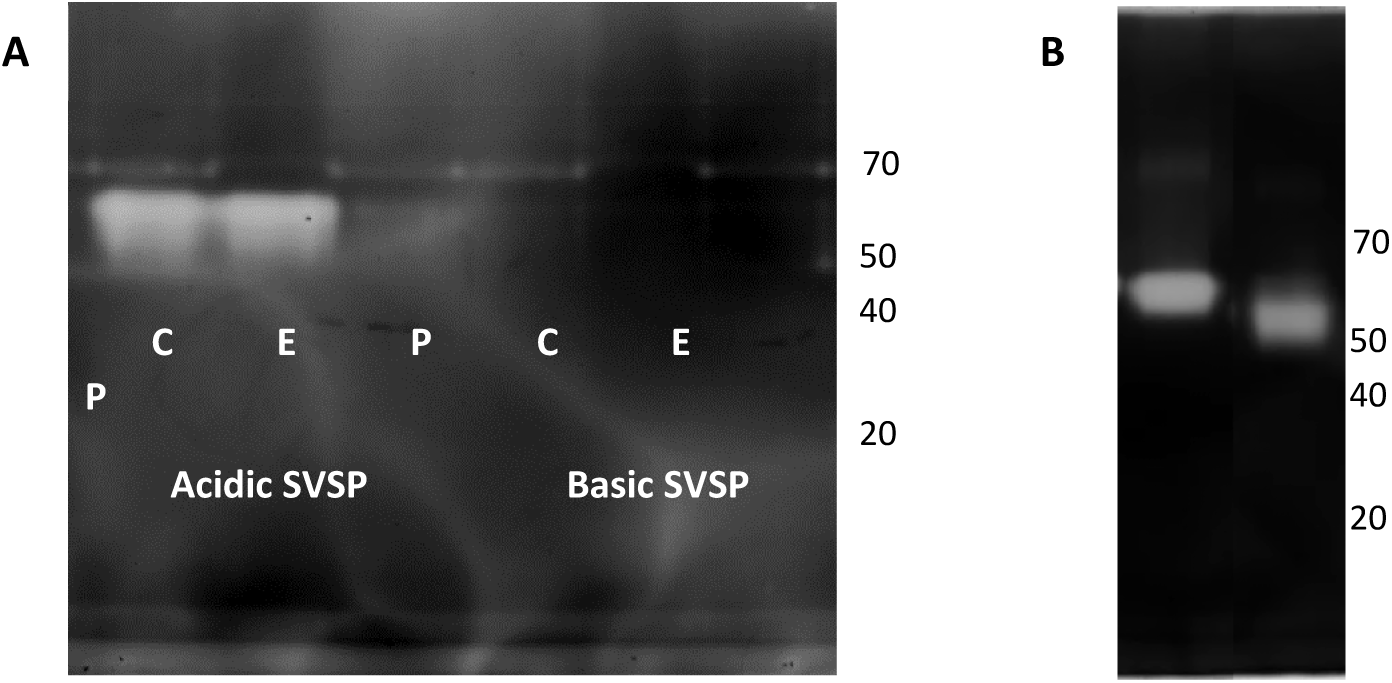
Gelatinase activity (gelatin zymograms) of the SVSPs from NGA *B. arietans* venoms. The zymograms were prepared with 0.2% (w/w) gelatin in a 10% acrylamide SDS-PAGE gel, which was run under non-reducing conditions. The zymograms were then incubated as in section 2.4.4.1 and then visualised by staining with Coomassie Blue R250. **A**: purified acidic and basic proteases (0.1 μg) with inhibitors. C, control (no inhibitor), E, 10 mM EDTA; P, 2 mM PMSF. **B**. Protein (0.5 μg) from peak 1 and 2 of RP-HPLC of acidic SVSPs (Fig. 3C). Molecular weight markers (Thermo Page Ruler) are indicated in kDa on the right of each zymogram.

An attempt was made to determine the importance of the N-glycans to the gelatinase activity (Fig. S5). Deglycosylation was carried out under native conditions, but this resulted in incomplete removal of the N-glycans (Fig. S5A). The partially deglycosylated material appears to have lost no gelatinase activity, however (Fig. S5B).

#### 3.3.2. Fibrinogen and prothrombin degradation

The purified SVSPs were tested for proteolytic activity against the key plasma proteins commonly targeted by haemotoxic venom proteases: fibrinogen and prothrombin (Fig. 7). The NGA basic and acidic SVSPs used for this analysis were the proteins in the main ion exchange peaks (Figs. 2A and 3A). Both acidic and basic SVSPs were able to digest fibrinogen to some degree but the degradation pattern for the acidic SVSP (lane A1) differs from that of the basic (lane B2). The acidic SVSP showed limited cleavage of both the alpha band, to a position just above the beta band in the control lane, and the beta band to the same position as the gamma band. The basic SVSP appears to possess alpha-fibrinogenase activity, with no apparent degradation of the beta and gamma bands. Neither form showed any proteolytic activity against prothrombin (Fig. 7B).

**Fig. 7.**
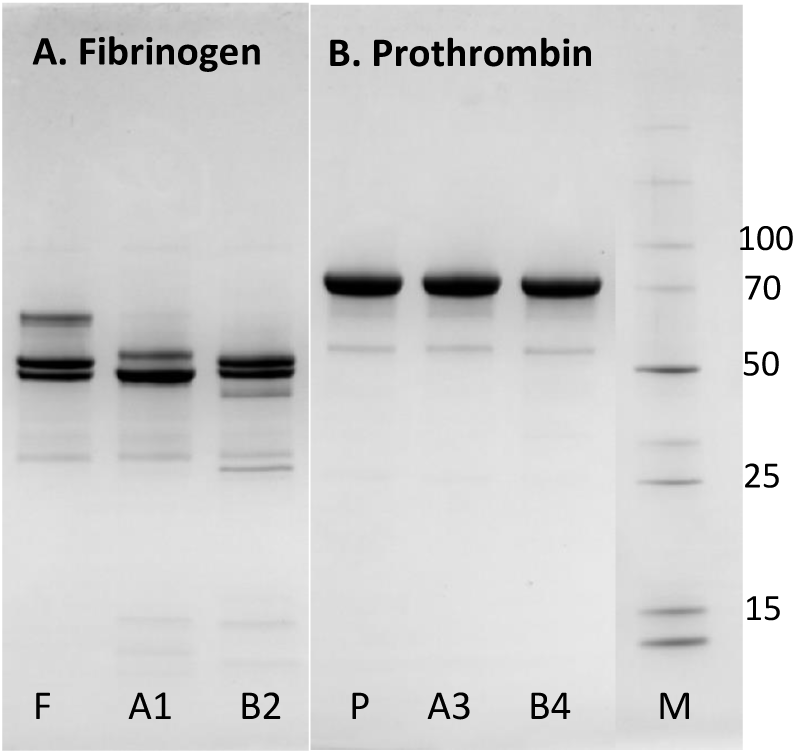
Degradation of fibrinogen and prothrombin by purified *B. arietans* serine proteases. Fibrinogen (gel A) or prothrombin (gel B) were incubated at 37°C for 120 minutes with the proteases at a 30:1 (w/w) ratio of substrate:protease. The gel was run under reducing conditions and the equivalent of 1.0 μg of substrate protein (prothrombin, fibrinogen) was loaded per lane. The gel is 4-20% acrylamide (BioRad) and was stained using Coomassie Blue R250. F is fibrinogen control and P is the same for prothrombin (lane 1 on both gels). Lane A1, F + acidic SVSP; lane B2, F + basic SVSP, lane A3, P + acidic SVSP; lane B4, P + basic SVSP. The molecular weights of key markers (lane M, Promega Broad Range) are indicated in kDa.

#### 3.3.3. Basement membrane protein degradation

As was the case for prothrombin, the serine proteases had no significant effect on basement membrane proteins (Fig.8A, lanes 1 and 2).

**Fig. 8.**
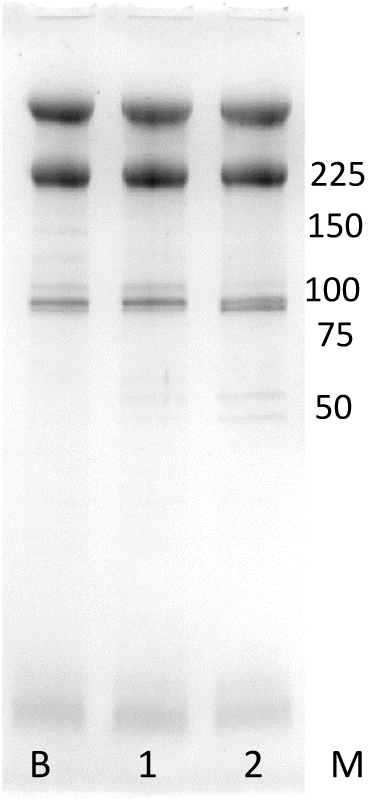
Degradation of basement membrane proteins by serine proteases isolated from *B. arietans* venoms. A basement membrane protein extract (Geltrex) was incubated at 37°C for 30 minutes with the purified proteases at a 30:1 (w/w) ratio of substrate:protease. The gel was run under reducing conditions and the equivalent of 8 μg of substrate protein was loaded per lane. The gel is 4-20% acrylamide (BioRad) and was stained using Coomassie Blue R250. Lane B, basement membrane protein control; lane 1, NGA acidic SVSP; lane 2, NGA basic SVSP. The molecular weights of key markers (lane M, Promega Broad Range) are indicated in kDa.

## 4. Discussion

A complete set of SVSPs was isolated from Nigerian puff adder venoms which had previously been shown to be rich in this class of protease. During this procedure, these SVSPs separated into two groups: basic and acidic forms. Despite their similarity in molecular weights and in their degree of N-glycosylation, each group possessed distinct enzymatic activities towards a variety of protein substrates. Using small chromogenic substrates, the acidic SVSPs were shown to be trypsin like, but the basic SVSPs were more chymotrypsin like. A bioinformatic analysis of SVSP sequences in a Nigerian puff adder transcriptome supported these findings and this will be expanded upon in this section.

Differences were observed between the ability of the two forms to degrade the general protease substrates casein and insulin B chain, with the basic form apparently acting with more potency and a broader specificity. Both forms of SVSP were able to degrade casein, but with clear differences in the extent: basic SVSPs fully degraded both casein subunits under the conditions used, but the acidic forms were only able degrade the upper band (α subunit). This difference in activity was more pronounced with insulin B chain: basic SVSPs cleaved the latter at multiple sites, but the acidic forms were unable to cleave any of the bonds this peptide. Both cleaved fibrinogen in a manner typical of SVSPs, but their precise actions were different. The acidic SVSPs cleaved both alpha and beta fibrinogen in a pattern that could be described as thrombin-like, as might be expected for a trypsin-like SVSP, whereas the basic SVSPs demonstrated alpha-only fibrinogenase activity.

The most interesting difference in the activities of the two forms of SVSPs was in their actions against gelatin. The acidic SVSPs were found to be responsible for the strong gelatinase activity observed in NGA venoms, whereas the basic SVSPs possessed no such activity. Gelatinase activity in venoms is usually associated with the SVMPs and such activity has not been previously determined in an isolated SVSP, which up to now have been shown to act only on plasma proteins (see Serrano, 2013). There is evidence in the literature for such activity, however: a PMSF-inhibited 36 kDa gelatinase was found on zymograms of *B. nasicornis* whole venom (Paixão-Cavalcante et al., 2015). Such activity, at around 50-60 kDa, was also observed in 11 of 12 NGA *B. arietans* tested venoms in an earlier study (Currier et al., 2010), but was reasonably assumed to be due to the action of an SVMP, since such activity would be unexpected for a SVSP based on the knowledge at the time. This activity may be peculiar to *Bitis* sp.

The availability of a full set of transcriptome data for Nigerian and Tanzanian snakes allowed us to match the biochemical characteristics of the purified SVSPs to predicted primary structures. An analysis of the NGA puff adder transcriptome found it to contain 24 different SVSP sequences, but only 6 were found in that of TZA snakes, mirroring the quantitative results of Dawson et al. (2025). Within this Nigerian set there is a broad range of predicted isoelectric points, ranging from 5.1 to 8.7 (not including contribution from sialic acids of the N-glycans) but a full analysis of the sequences indicated the presence of two distinct groups of acidic and basic proteases. The numbers of predicted N-glycans also matched the experimentally determined values: all the basic forms are predicted to have 4 or 5 N-glycans and the acidic forms 5 or 6 N-glycans, except for one acidic form with just 3 predicted N-glycan sites. Critical differences can be seen at their respective substrate-binding sites, however. Figure 9 shows the relevant sections of the four NGA *B. arietans* SVSP transcripts which matched to the SVSPs isolated here.

**Fig. 9.**
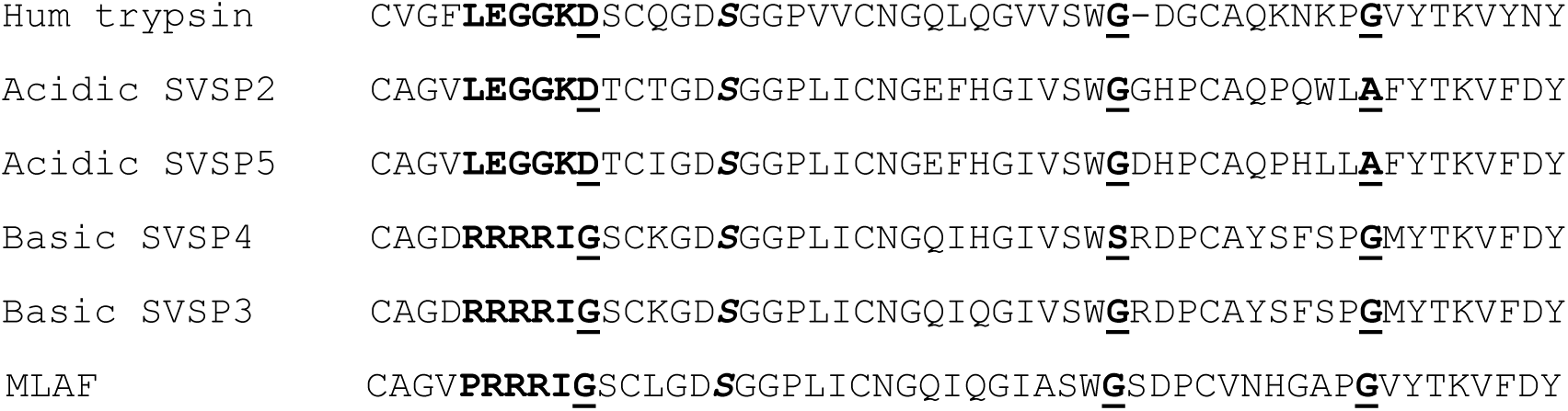
Sequence alignment of the substrate-binding region of the most abundant acidic and basic SVSP transcripts from NGA puff adder. The four NGA SVSPs sequences (see Fig. 5) were subject to a CLUSTAL multiple sequence alignment by MUSCLE 3.8 (Larkin et al., 2007). Also included for comparison are sequences from human trypsin-1 (P07477) and *M. lebetina* alpha-fibrinogenase MLAF (Q8JH85 VSPA_MACLB). In bold is the S_1_ residue (D189 or G189) plus the 5 residues immediately N-terminal to this. The active site S195 is shown in bold/italics. Shown in bold/underlined are the three key substrate pocket residues 189, 216 and 226 (DGG in trypsin; DGA in the acidic SVSPs; GGG or GSG in basic SVSPs and GGG in MLAF). All residues are numbered using the chymotrypsin system.

The transcriptome data for the NGA acidic SVSPs support the experimental findings (Table 1) in that they are trypsin-like serine proteases, having an aspartate (**<u>D</u>** in Fig. 9) at the bottom of the substrate pocket (S_1_ binding site, residue 189 using chymotrypsin numbering). They also have sequences immediately N-terminal to the substrate-binding site (VLEGGKDT) similar to those of other well-studied SVSPs, such as batroxobin (Itoh et al., 1987). Thus, in most respects, these *B. arietans* acidic SVSPs are typical snake venom serine proteases (Serrano, 2013), though no others have so far been shown to have the gelatinase activity of the acidic SVSPs isolated here. At the two other positions that form the substrate pocket, 216 and 226, the acidic SVSPs have glycine (G) and alanine (A) residues respectively. The alanine at position 226 in lieu of the smaller glycine residue found here in trypsin and thrombin might restrict access of large residues in the P_I_ position of the substrate. Despite this, however, it appears to cleave both alpha and beta fibrinogen in a similar manner to thrombin (Fig. 7A), which is quite rare for an SVSP (Kini, 2005).

It is difficult to explain the inability of the trypsin-like acidic SVSP to cleave at the lysine (K) and arginine (R) residues in insulin B. Nor did it show any cleavage of angiotensin I, in which the second residue is an R. Despite their being different classes of protease, it is interesting that both the puff adder SVMP PIII-c gelatinase isolated previously (Wilkinson et al. 2025) and acidic SVSP gelatinase described here are unable to degrade insulin B, which is routinely used as a venom protease substrate because of its wide variety of peptide bonds. We have isolated a gelatinase SVMP PIII-a from *Echis romani* venom which is also inactive towards insulin B, in contrast to all the other (non-gelatinolytic) SVMP PIIIs isolated from the same venom (manuscript in preparation). This all suggests that gelatinase activity in venom proteases is quite specific, perhaps with a primary specificity to amino acids at both P_1_ and P’_1_. This may point to a genuine functional role and it not being simply a case of a protease acting against a general protein substrate, such as is the case for other commonly used protease substrates, e.g. casein. Because of the possible functional and clinical relevance of the ability of a venom protease to degrade gelatin (predominantly denatured collagen I, which is rich in Lys and Arg residues) these SVSP and SVMP gelatinases warrant further studies beyond that of a simple scientific interest.

In contrast to the acidic SVSPs, the basic forms have a Gly in lieu of the Asp at position 189, contained within a distinct arginine-rich sequence (DRRRRIG) on the N-terminal side of the substrate-binding site. This is partly responsible for their basic pI and is found in very few SVSPs: a BLAST search found just three such proteins: rhinocerases (Vaiyapuri et al., 2011), alpha-fibrinogenase ML-AF (Siigur, 2003) and an *E. ocellatus* serine protease (submitted to GenBank: ADE45139.1). The G189 at S_1_ is the basis of the non-trypsin-like activity of these basic SVSPs. They also have a Gly at positions 216 and 226, and consequently a large G189G216G226 substrate-binding pocket which would explain the preference for phenylalanine at P_1_ (Table 1). This is similar to the action of chymotrypsin which has a large (S189G216G226) substrate pocket, but the specific activity of the basic SVSP towards the Phe-containing nitroanilide was far lower than that of chymotrypsin. This NGA basic SVSP is able to cleave insulin B at multiple sites (Fig. S4), more than might be expected for a peptide substrate with three phenylalanine residues, two of which are adjacent, so it must be able to cleave at other amino acid residues. Insulin B chain possesses two His residues and this basic SVSP was able to convert angiotensin I to II (Fig. S6), requiring cleavage of a His-Pro bond. Other SVSPs have been predicted to have the same G189G216G226 substrate pocket (Vaiyapuri et al., 2012), but the only one that has been fully characterised is the alpha fibrinogenase ML-AF of *M. lebetina* (Siigur et al.,, 2003) and as well as having the same fibrinogen specificity as the puff adder basic SVSP, it also displays the same inactivity towards Arg- and Tyr-esters (Samel et al., 2002). ML-AF also has the arginine-rich sequence (PRRRRIG) and is a basic SVSP (see Fig. 9). It also shows a preference for phenylalanine, cleaving insulin B at the Phe-Phe and Phe-Tyr peptide bonds, but also at Tyr-Leu (Mahar et al., 1987). Thus it seems that ML-AF, described as unique in Siigur et al., (2003), and the puff adder basic SVSP are related proteins. Alpha fibrinogen degradation may be their principal role, contributing to an anticoagulant action of the respective venoms.

SVSPs are generally placed into either of two classes (Serrano and Maroun, 2005): acidic forms which act against plasma proteins involved in haemostasis, and basic SVSPs with direct platelet-activating (PA) activities. The puff adder acidic SVSPs appear to conform to this sub-classification, but the basic SVSPs do not. The known basic PA-SVSPs (e.g. PA-BJ (Serrano et al., 1995) and the thrombocytins (Niewiarowski et al., 1979; Hill-Eubanks et al. 1989)) are far more basic (pIs >8) and a lot smaller (28 kDa with just one N-glycan) than the puff adder basic SVSP. But critically, the PA-SVSPs are trypsin-like, exerting their action through cleavage at Arg/Lys-xxx peptide bonds.

### Potential pathology implications

The main purpose of this work was to expand our knowledge of the puff adder SVSPs which were identified to be a quantitatively significant component of the venom in previous studies. Much has been revealed concerning their biochemistry, but to extrapolate the results of the assays to possible pathological effects can only be done at this stage by comparison with what is known from studies on other venom proteases. The ability of the gelatinolytic acidic SVSPs to degrade collagen I, the major form of collagen in mammalian tissues, suggests it may play a role in many of the clinical features of puff adder bites that might arise as a consequence of tissue damage. These would include damage at the bite site, often leading to necrosis, but also blisters which commonly form distal from the bite (Warrell et al., 1975; Tianyi et al., 2025). It has been proposed that snakebite-induced blistering can arise due to damage to fibrillar collagens at the dermal-epidermal junctions (Gutiérrez et al., 2018; Jiménez et al., 2008) and the gelatinolytic puff adder SVSPs may be responsible for this.

Having the ability to degrade fibrinogens, both forms of SVSP will undoubtedly contribute to the anticoagulant outcome of puff adder envenomation by affecting the clottability of fibrinogen by thrombin. Such a role for SVSPs is well documented (Serrano and Maroun, 2005; Kini, 2005), but in addition to this, the ability of the acidic forms to degrade collagen may also exacerbate the anticoagulant effects of the venom, since this protein plays a key role in the initial adherence of platelets to damaged blood vessel wall (Jackson, 2007).

The possible role of the SVSPs in producing the hypotension that is commonly seen in puff adder bite victims has not been studied in detail here. The basic SVSP was able to convert angiotensin I to II (Fig. S6). Angiotensin II is vasoconstrictive rather than vasodilatory, but this result would need to be placed in context of the action of whole venom on angiotensin I; the SVSP action is very likely to be overridden by the action of the highly potent puff adder SVMP PIs. Amongst the various SVSP proteins isolated here are likely to be the kallikrein-like *B. arietans* SVSPs that have previously been characterised as having the ability to generate kinins which will act in a vasodilatory manner. Within the acidic SVSP transcripts are sequences that match with the N-terminal sequence that Nikai et al. (1993) found for their 58 kDa acidic SVSP, and the tryptic peptide sequences from Kn-Ba obtained by Megale et al. (2018) matched exactly with the one of the NGA acidic SVSP transcripts (NIG_SVSP_7 Contig4601||Toxin2176, see Fig. S7). Kn-Ba is only 33 kDa in size, but the sequence of SVSP-7 predicts just 3 N-glycans (as opposed to the 5 or 6 predicted for the other 24 transcripts), so it would be considerably smaller than the main group of SVSPs. This may be the small acidic SVSP in lane 1 of Fig. 3B.

## 5. Conclusion

This study clearly identifies the protein responsible for the strong gelatinase activity we previously detected whilst isolating and characterising puff adder SVMPs (Wilkinson et al., 2025). The activity is due to an acidic serine protease and is a novel activity for an SVSP, an activity usually associate with SVMPs. At least as important as this biochemical peculiarity is the observation that the activity is highly variable geographically. This variety will further compound the issues discussed in the SVMP paper (Wilkinson et al., 2025). In that study we identified two forms of an SVMP PI (arilysin), one of which was much more destructive towards protein substrates, particularly ECM components, than the other. The presence of these two SVMPs in puff adder venoms also varied geographically. In the case of the two main venoms we have studied, the Nigerian had the weaker arilysin, but has the gelatinolytic SVSP. In contrast the Tanzanian venoms had the potent arilysin, but no gelatinolytic activity. Others, e.g. Ghana and Eswatini had both the potent arilysin and the gelatinase SVSP, and in the previous study (Wilkinson et al., 2025) the gelatinase activity in Kenyan venoms (with the weaker arilysin) was due to an SVMP PIII-c rather than the SVSP. Since these bioactivities will play a major role in snakebite pathology, it is not surprising, therefore, that the clinical features associated with puff adder envenomation are greatly variable from bite to bite. And it is clearly evident from these two studies that development of a pan-African therapeutic for puff adder snakebite will need to take into account this geographical variation in the key venom components and their relative toxicity.

## Acknowledgements

We thank Herpetologists Paul Rowley and Edouard Crittenden at LSTM for the maintenance and husbandry of the snake collection and the provision of venom samples.

The authors wish to thank the following for providing financial support to this study: Wellcome Trust grant 223619/Z/21/Z (to M.C.W., C.M.M., R.A.H., N.R.C.), Wellcome Trust grant 221712/Z/20/Z (to R.A.H., N.R.C.), The UK Foreign Commonwealth & Development Office grant 300341-115 (to R.A.H., N.R.C.), Department of Defense grant ID07200010-301-35 (to A.S.).

## Supplemental figures [Wilkinson et al 2025b]

**Fig. S1.**
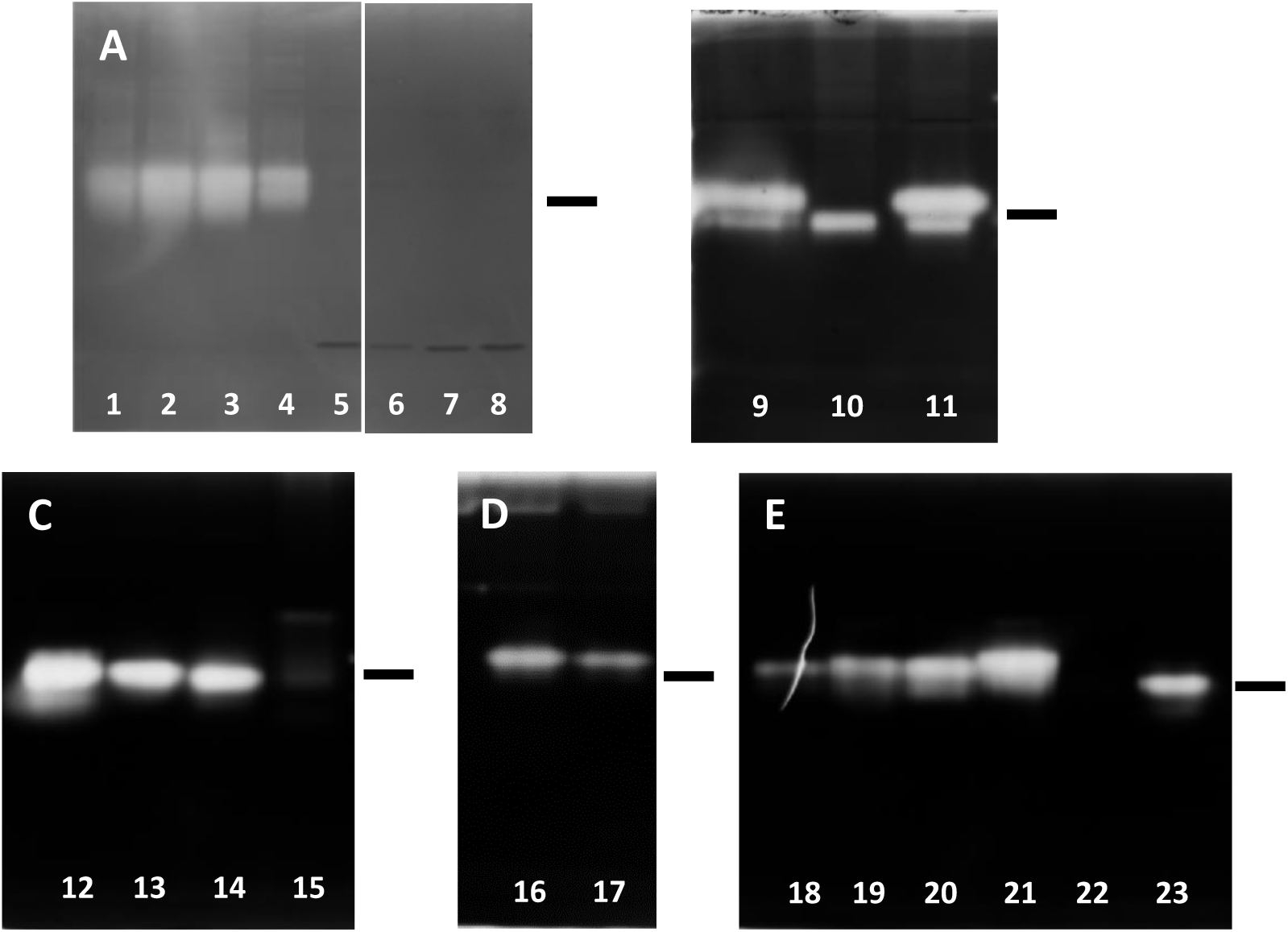
Gelatinase activity (gelatin zymogram) of venoms taken from various African and Saudi Arabian puff adders. The zymograms were prepared with gelatin included at 0.1% (Gel A, supplied by Thermo-Fisher) or 0.2% (Gels B-E, made in-house) in a 10% acrylamide SDS-PAGE gel. The gels were run under non-reducing conditions and 2 μg of venom was loaded per lane. The zymograms were then incubated as in section 2.4.4.1 and then visualised by staining with Coomassie Blue R250. **Gel A:** Venoms from snakes held at LSTM at the time of the study. Lanes 1 - 4: NGA and lanes 5-8: TZA. **Gel B:** Lanes 9 – 11, venoms from three captive bred Kenyan snakes (LSTM). **Gel C:** Lane 12, Venom from LSTM Eswatini snake; lane 13, stock Zimbabwe venom; lanes 14 and 15, stock South African venoms. **Gel D:** lanes 16, 17, venoms from two Ghanaian snakes formerly housed at LSTM. **Gel E:** Lanes 18 - 21 venoms from four Nigerian snakes formerly housed at LSTM; lane 22 Saudi Arabian stock venom; lane 23, Malawi Stock venom. The bar to the right of each gel indicates the position of the 50 kDa molecular weight marker, revealed following complete staining of the gel.

**Fig. S2.**
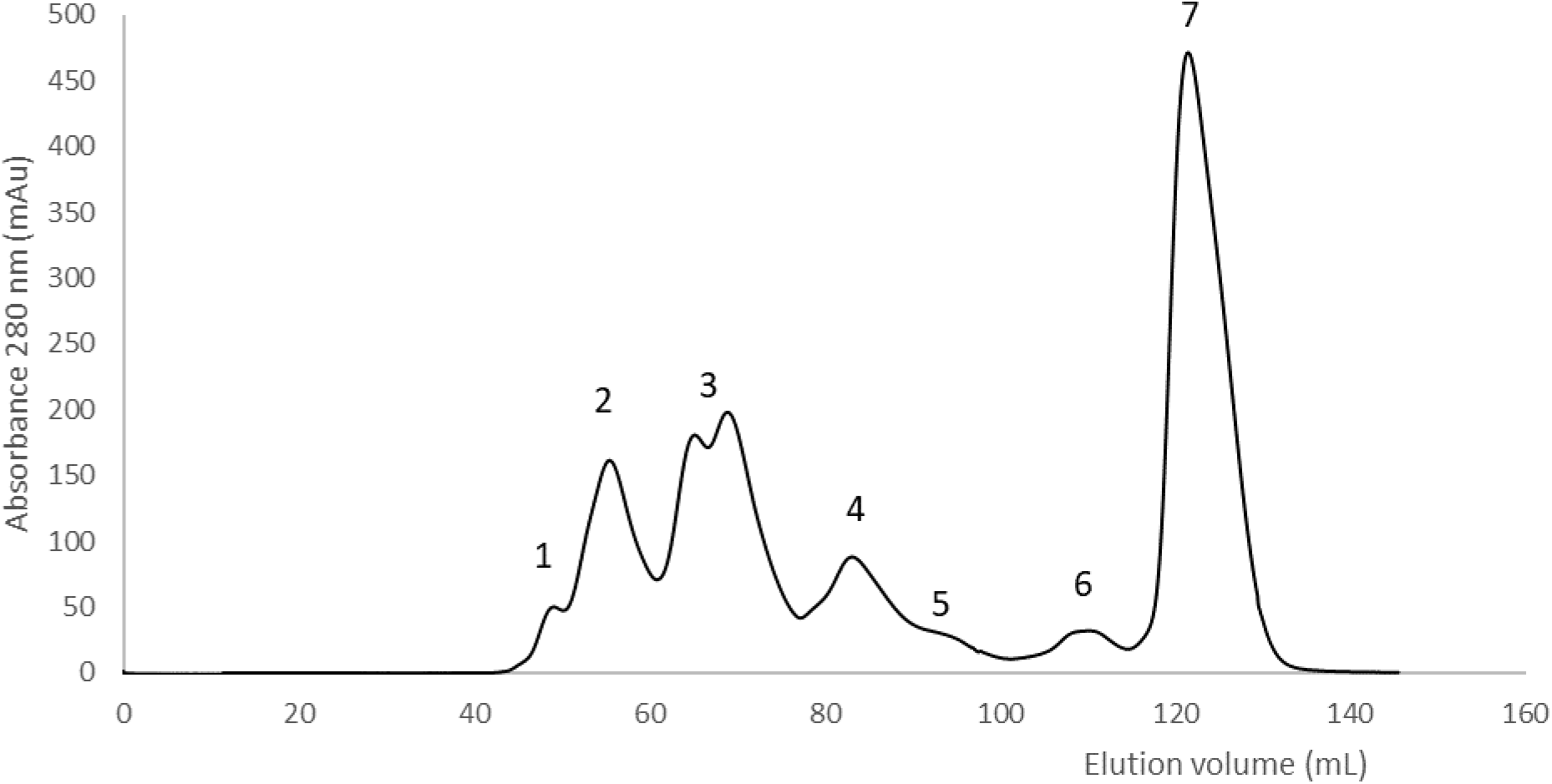
Size exclusion chromatography separation of NGA *B. arietans* venom. Whole venom (20 mg NGA 011) was separated on a 120 mL column of Superdex 200HR equilibrated in 50 mM sodium phosphate pH 5.2. The flow-rate was 1.0 mL/min and elution was monitored at 280 nm

**Fig. S3.**
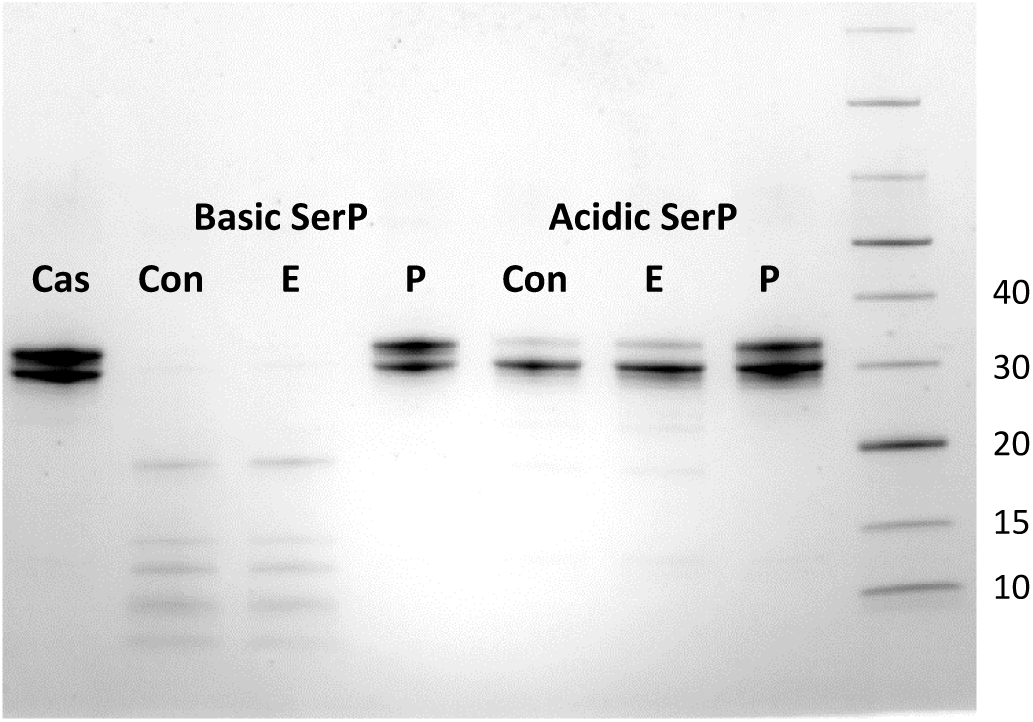
Casein degradation assay: Purified serine proteases from NGA puff adder. Basic or acidic SerPs from the ion exchange chromatography steps were incubated with bovine casein, either alone or with 5 mM EDTA or 2 mM PMSF, for 90 minutes at 37°C. An aliquot of the reaction mix equivalent to 2 ug of casein was run on SDS-PAGE. The gel used was 4-20% acrylamide (BioRad) and was stained with Coomassie Blue R250. Cas, casein control; Con, SVSP with casein; E, SVSP + casein + 5 mM EDTA; P, SVSP + casein + 2 mM PMSF. Molecular weight markers (Thermo Page Ruler) are indicated in kDa on the right.

**Fig. S4.**
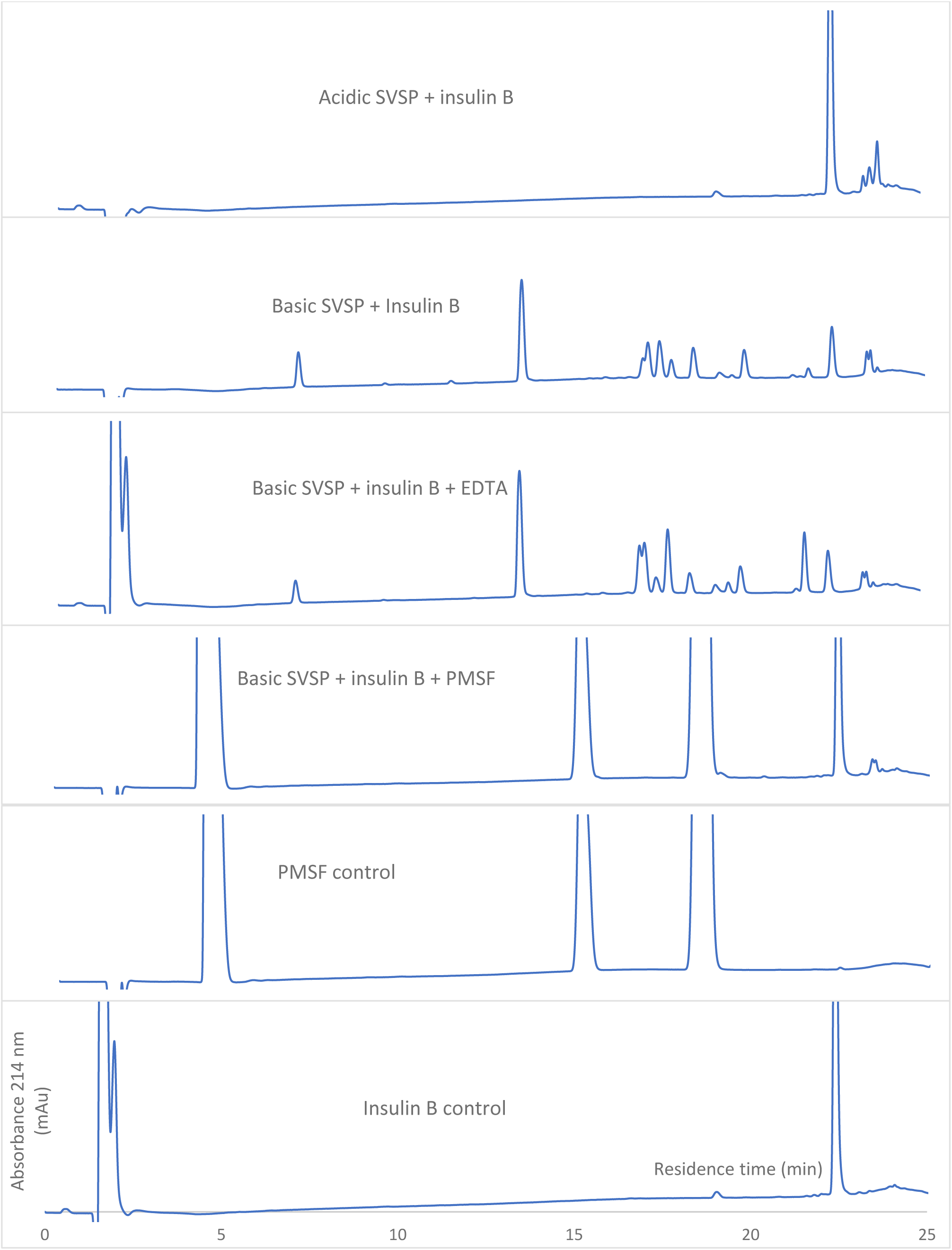
Insulin B degradation assay: Purified SVSPs. The main acidic and basic SVSPs from the ion exchange chromatography steps were incubated at 37°C for 90 mins with insulin B chain at a ratio of 30:1 (w/w) insulin B: SVSP. An aliquot containing the equivalent of 1 ug insulin B was analysed by RP-HPLC with monitoring at 214 nm. The conditions for each assay are indicated in the inset.

**Fig. S5.**
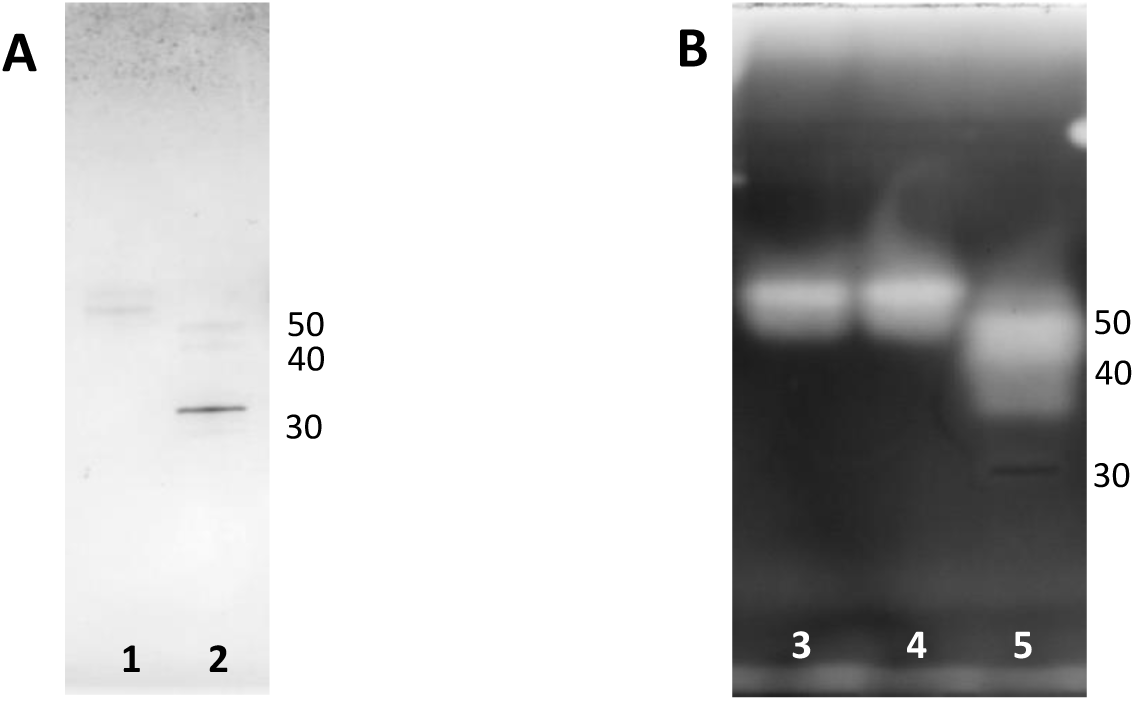
Effect of deglycosylation on the gelatinase activity of the acidic SVSP. The protein used was 0.2 μg of peak 1 of RP-HPLC of acidic SVSPs (Fig. 3C). **A**: Standard reduced SDS-PAGE analysis of deglycosylated acidic SVSP. Lane 1 control, no PNGase F; lane 2 PNGase F (visible at 32 kDa) treated. **B**. Gelatin zymogram, lanes 3, untreated SVSP; lane 4, control, no PNGase F; lane 5 PNGase F treated. The zymogram was prepared and run as in previous experiments then visualised by staining with Coomassie Blue R250. Molecular weight markers are indicated in kDa on the right of each zymogram.

**Fig. S6.**
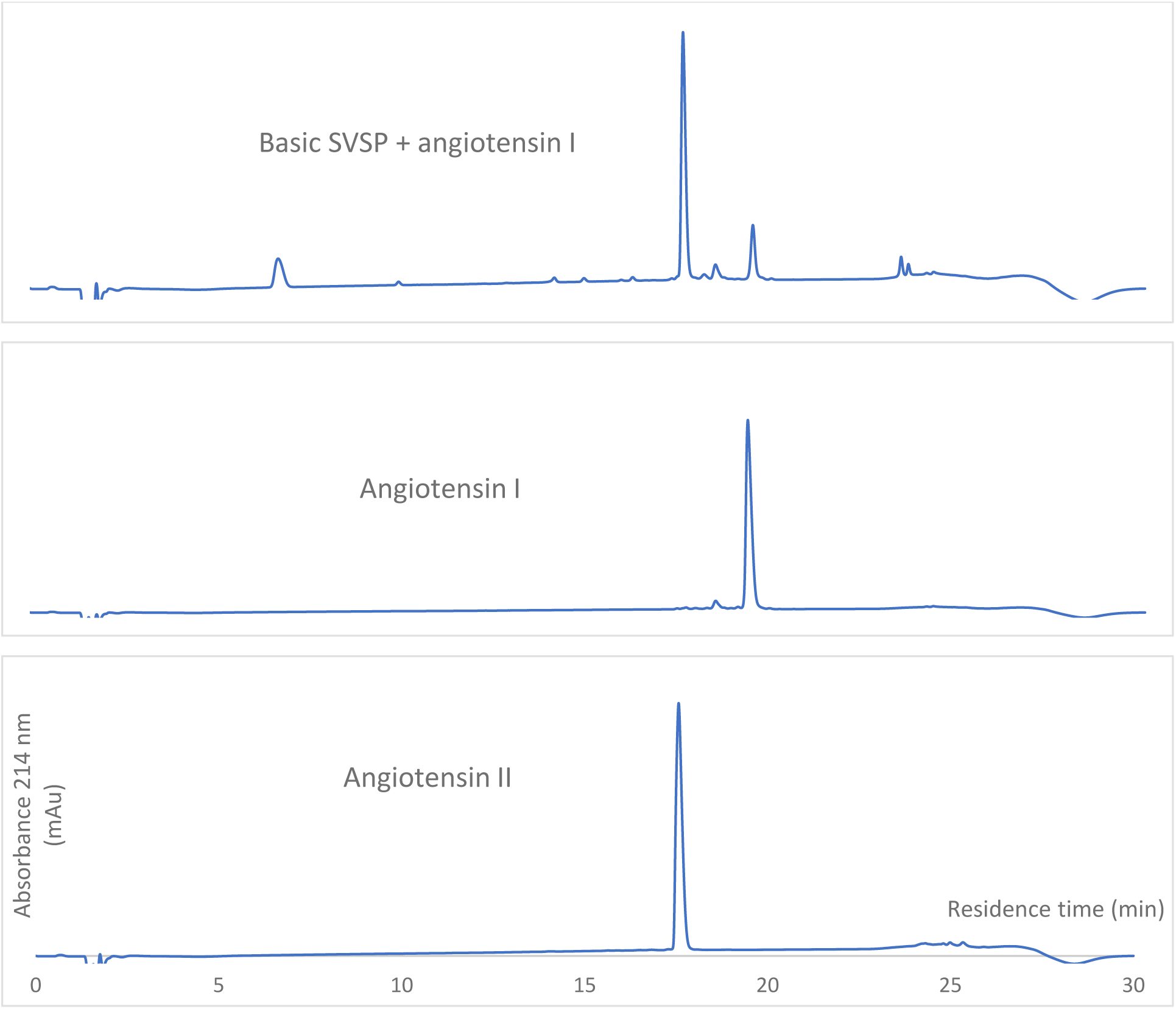
Action of the basic SVSP on angiotensin. **I.** The main basic SVSPs from the ion exchange chromatography steps were incubated at 37°C for 60 mins with angiotensin I at a ratio of 30:1 (w/w) angiotensin I:SVSP. An aliquot containing the equivalent of 1 ug angiotensin I was analysed by RP-HPLC with monitoring at 214 nm (top panel). The lower panels show angiotensin I and II standards.

**Fig. S7.**
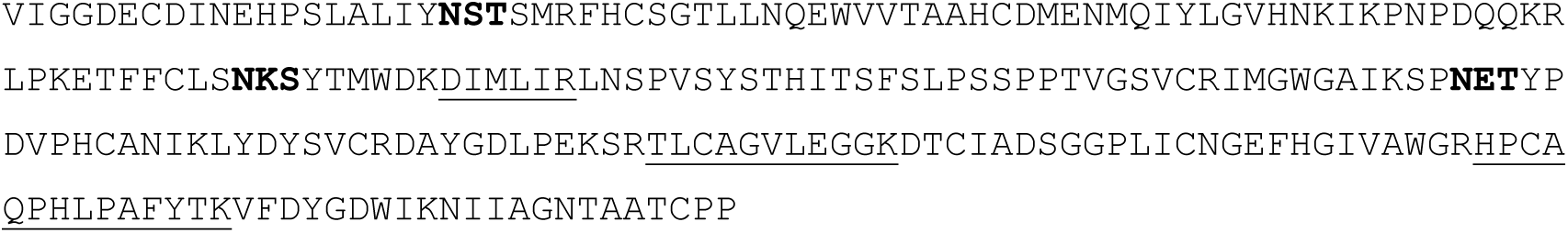
Amino acid sequence for acidic SVSP transcript NIG_SVSP_7 Contig4601||Toxin2176. This SVSP was unusual within the NGA SVSP transcripts in that it possesses only 3 predicted N-glycan sites (bold). It has tryptic peptides (underlined) to exactly match those found in the 33 kDa kinin-releasing SVSP Kn-Ba isolated by Megale et al. (2018)

